# Influencing factors of telephone-cardiopulmonary resuscitation in China: a qualitative exploration based on managerial perspectives

**DOI:** 10.1101/775874

**Authors:** Xuehua Zhu, Li Gui, Ying Chen, Yin Lin

## Abstract

**Background:** Telephone-cardiopulmonary resuscitation(T-CPR) has been proven to systematically improve bystander CPR implementation and thus improve the survival rate of out-of-hospital cardiac arrest (OHCA) patients on a large scale. However, China has a lower proportion of cities that provide T-CPR than other countries.This study aimed to explore the factors affecting the providing of T-CPR based on managerial perspectives and promote the implementation of T-CPR in China to Protect human health.

**Methods:** This study adopted a descriptive qualitative method.The managers from health bureau and first-aid Center were recruited to participate through purposive sampling. Data were collected using semi-structured interviews and Colaizzi 7-step analysis method was adopted to summarize and conclude the theme.

**Results:** A total of 10 managers were interviewed.Five main themes were identified: (a) bystander factors, (b) dispatching factors, (c) legal factors, (d) guiding factors, and (e) financial factors.

**Conclusion:** It is urgent to promote the implementation of T-CPR in China.We can promote it by strengthening the training of bystanders in CPR knowledge and skills, developing T-CPR guidance process suitable for Chines national conditions, building an intelligent prehospital emergency system, promoting the legislation of first aid exemption, and providing financial support from various channels.

## 1 INTRODUCTION

Out-of-hospital cardiac arrest (OHCA) is the most urgent and dangerous public health issue in the world (Benjamin et al.,2017;Miller & Falk,2020;Høybye et al., 2021).Modern medicine has fully proved that 4 to 6 minutes after cardiac arrest is the golden time for rescue, but it is difficult for ambulances to arrive at the scene in the golden time. Therefore, whether the public implement cardiopulmonary resuscitation (CPR) or not and the quality of CPR have a significant impact on the survival rate of OHCA patients (Sutter et al.,2015).Because the witnesses are often non-professional medical personnel, they will not really administer effective CPR for patients with cardiac arrest because they are not confident in their own CPR skills, afraid of the implementation of mouth-to-mouth artificial respiratory infectious diseases, panic, and have not learned CPR (Becker et al.,2019;Hagihara et al.,2018).So that patients miss the best time to rescue and lead to serious consequences, bringing heavy burdens to society and families.

In order to increase bystander cardiopulmonary resuscitation (B-CPR) and to improve patient outcomes, American heart association (AHA) Guidelines for Cardiopulmonary Resuscitation and Emergency Cardiovascular Care (ECC) clearly point out that the first witnesses who have never received CPR training or who have received training but failed to successfully initiate CPR in the first time should be instructed by dispatchers through telephone to implement CPR (T-CPR) (Dobbie et al.,2018).T-CPR enables some first-time witnesses who have never received CPR training to implement CPR under the guidance of the dispatcher’s telephone, thus making it possible for more OHCA patients receive early CPR.

Since telephone-assisted cardiopulmonary resuscitation can significantly improve the willingness and quality of witnesses to perform CPR, T-CPR has been carried out in many countries, such as the United States, Britain, Japan, Sweden, Canada, Switzerland and Norway(Kleinman et al.,2015;Bobrow et al.,2016;Shimamoto et al.,2015).But T-CPR is seldom carried out in China, only in a few cities such as Suzhou and Foshan.China, a country with 4/1 of the world’s population, in order to improve its popularity of T-CPR and the success rate of on-site rescue, and to reduce the mortality and disability rate after cardiac arrest, our study is based on managerial perspectives (Health Bureau, Fist-aid Center and other management personnel) who will play a key role in carrying out TCPR, and through a qualitative study to explore the factors affecting the providing of TCPR, especially finding out the possible obstacles, so as to promote the implementation of TCPR and play a role in promoting human health.In addition,it can effectively solve the occupational problems related to the field of pre-hospital emergency care to improve the occupational environment of occupational health care workers.

## 2 METHODS

Our main aim was to identify influencing factors of telephone-cardiopulmonary resuscitation in China based on managerial perspectives.

### 2.1 Participants

A purposive sampling approach, was used to select the managers were recruited from health bureau and first-aid Center.The sample size is based on the principle of information saturation.Finally, a total of 10 managers consented to participate in the study. The inclusion criteria were: (a) managers from health bureau and first-aid Center aged 18-60, (b) worked in this management for more than 2 years, (c) could understand and communicate in Chinese, (d) agreed with the interviewees. The exclusion criteria were: (a) mental disorders and/or cognitive impairment, (b) unable to complete the study due to physical condition, (c) unable to perform CPR due to physical disability or other reasons.

### 2.2 Study design and methodology

The study adopted a descriptive qualitative method.Data were collected by using semi-structured interviews through face-to-face or by telephone. The interview outline was designed according to the previous literature review and research purposes.The final interview guide was mainly included the following contents: (a) demographic information;(b) views and opinion on performing on-site cardiopulmonary resuscitation (CPR) and telephone-assisted cardiopulmonary resuscitation (T-CPR) for the public; (c) the factors affect the public and regulatory acceptance of T-CPR in China; (d) the factors or difficulties affect the implementation of T-CPR in China based on managerial perspectives; and (e) suggestions to promote carry out T-CPR in China.

### 2.3 Data collection

The face-to-face interviews lasted from 20-50 minutes (mean±SD: 32.5) and were conducted in a location chosen by the participants.We collected data by means of a semi-structured interview, and all sessions were audio-recorded.At the beginning of the interviews, the researcher introduced herself and explained the purpose and significance of this study to the participants. After obtaining the formal consent of them, the informed consent was signed. The researcher remained neutral and avoided making any judgment on the interview contents.The privacies of the participants were protected, and the name was replaced by the code.

### 2.4 Data analysis

After one interview was completed, the recording was transcribed verbatim into text within 24 hours. The data were analyzed concurrently with the data collection using the Colaizzi 7-step analysis method. The first step involved reading the full interview transcript several times to have a comprehensive understanding of the participant’s experiences. Then, any narrative data related to the impact of the T-CPR implementation were hand-coded line-by-line. Next, generated the theme based on the similarities and differences between the code and the concept. Finally, created a definition for each theme and selected the support references from the data. The data collection and data analysis procedures were conducted repeatedly until there was no more topic (Glenn & Bowen,2008).

### 2.5 Ethical considerations

We obtained approval by the ethical committee of the institutions(20200529-1) for this study. All participants were recruited as volunteers. Before the interviews, participants were given a participant information sheet with written information on the project and provided informed consent. All reports were kept private and confidential. Code numbers were used in this article to identify individual comments.

### 2.6 Rigour

Four criteria, including credibility, dependability, conform ability, and transfer ability, were used to evaluate the rigor of the qualitative research.Interviews were audio-recorded to ensure credibility and to ensure that participants’ responses were captured accurately. In addition, participants’ statements were probed and clarified to ensure that they were understood accurately. An audit trail based on the interview outline, audio-recordings, transcripts, field notes, and data analysis process was clearly elaborated. Dependability and conform ability were also established by having decisions relating to the study made by two other researchers in the team. Other than the primary researcher, two researchers who were experienced in the field of study were involved in data analysis.

## 3 RESULTS

Participants (n=10) were predominately male, Han Chinese, married, and between the ages of 40 and 50 years (see Table 1).Study results illuminated the influencing factors of telephone-cardiopulmonary resuscitation in China based on managerial perspectives.Transcription and coding uncovered five main themes: (a) bystander factors, (b) dispatching factors, (c) legal factors, (d) guiding factors, and (e) financial factors.

**Table 1.** Participant Demographics

| Characteristic | n | % |
| --- | --- | --- |
| Gender (n = 10) |  |  |
| Female | 2 | 20 |
| Male | 8 | 80 |
| Age (years)(n = 10) |  |  |
| 20 to 30 | 1 | 10 |
| 31 to 40 | 4 | 40 |
| 41 to 50 | 4 | 40 |
| 51 to 60 | 1 | 10 |
| Working time (years)(n = 10) |  |  |
| 1 to 10 | 3 | 30 |
| 11 to 20 | 2 | 20 |
| 21 to 30 | 5 | 50 |
| Education(n = 10) |  |  |
| Bachelor degree | 9 | 90 |
| Doctoral degree | 1 | 10 |
| Marital status(n = 10) |  |  |
| Never married | 1 | 10 |
| Married or with a partner | 9 | 90 |
| Profession title(n = 10) |  |  |
| Middle | 6 | 60 |
| Senior | 2 | 20 |
| Associate senior | 2 | 20 |
| Graduation major(n = 10) |  |  |
| Clinical medicine | 4 | 40 |
| Health management | 3 | 30 |
| Computer science | 2 | 20 |
| Preventive medicine | 1 | 10 |
| current work area(n = 10) |  |  |
| first-aid center | 5 | 50 |
| medical affairs under Provincial Department of Public Health | 1 | 10 |
| maternal and child Health Service | 1 | 10 |
| Red Cross ambulance training center | 2 | 20 |
| emergency office of the commission for health and discipline | 1 | 10 |
| position(n = 10) |  |  |
| Deputy director | 3 | 30 |
| Director | 4 | 40 |
| Chief | 3 | 30 |

### 3.1 Theme one:Bystander Factors

Bystander factors were used to code and categorize factors into those that (a) different knowledge levels of bystanders, (b) weak awareness of first aid and low level of cardiopulmonary resuscitation of bystanders and (c) volatile emotions of bystanders.

#### 3.1.1 Factors associated with different knowledge levels of bystanders

T-CPR implementation is influenced by the educational levels of bystanders.If the bystander’s education level is not high, it will reduce the effectiveness and timeliness of the dispatcher’s guidance.

> *“If he or she (refers to bystanders) has a higher level of knowledge, then our dispatcher’s guidance will be much better, and its effectiveness may be higher*.*In other words, the caller’s knowledge and education are related. If he or she doesn’t understand what we’re talking about, it may be difficult to implement T-CPR*.*” (P5)*

#### 3.1.2 Factors associated with weak awareness of first aid and low level of cardiopulmonary resuscitation of bystanders

At present, there is still a big gap in the awareness and level of first aid among Chinese residents compared with the developed countries. The national consciousness of first aid is so weak, and the basic knowledge of first aid is not valued. In a word,if bystanders have never been exposed to CPR learning, it is difficult to do so by telephone.

> *“Our country is, in my own city, uh (pause*…*), our common people have little knowledge of first aid, such as cardiopulmonary resuscitation. In this situation, the effect of T-CPR will be affected*.*To tell the truth, I think CPR is a little abstract, and it’s hard to make it clear on the phone. If someone hadn’t received any training in CPR, I don’t think he could implement TCPR*.*” (P7)*

#### 3.1.3 Factors associated with unstable emotions of bystanders

Bystanders will feel fear and dread when they encounter such sudden situations,which will affect the implementation of T-CPR:

> *“In the process of T-CPR, sometimes it is necessary to calm the caller constantly. In such a sudden situation, some callers are unstable and will be more anxious. Sometimes on the phone we will tell them what to do and how to do, but they don’t cooperate. They only ask you to send an ambulance as soon as possible. They don’t listen to the dispatcher at all. They don’t understand why they need to rescue the sudden cardiac arrest (SCA) patients*.*”(P7)*

### 3.2 Theme two:Dispatching Factors

Dispatching factors were categorized into those that associated with lack of professional skills of dispatchers, dispatching equipment,and standard procedures of dispatch.

#### 3.2.1 Factors associated with lack of professional skills of dispatchers

Because of the special nature of work, the particularity of working time, high recruitment requirements for recruiting, difficult promotion and low salary, the dispatchers are lacking and difficulty to recruit.Based on the difficulty of personnel recruitment, many first-aid centers can only reduce the recruitment requirements, so the first aid knowledge and skills of dispatchers in many dispatching posts are not high, which has a strong impact on the implementation of T-CPR in China:

> *“In our city, non-medical workers are answering the emergency calls now. They don’t have medical knowledge. If he or she makes a descriptive mistake, the consequence is that, as you know, it’s not necessarily good for the person’s rescue. He or she may be the defendant*.*” (P3)*

#### 3.2.2 Factors associated with lack of dispatching equipment

The investigation found that the current dispatching equipment was imported from abroad and not localized.In fact, it was not suitable for China in some cases. Therefore, the applicability of dispatching equipment affects the implementation of T-CPR.

> *“We need to establish a more standardized way of guiding. This medical priority dispatch system (MPDS) is relatively mature in the United States, but it needs to be localized if it is used in China. After all, MPDS is made in accordance with the national conditions of the United States*.*”(P2)*

#### 3.2.3 Factors associated with lack of standard procedures of dispatch

The lack of unified, simple and standardized flow chart to guide the dispatching process is also one of the important reasons that affect the implementation of T-CPR.

> *“We feel that such an approach (T-CPR) must be very meaningful. That is to say, the telephone guidance is very meaningful, but the key of T-CPR is to standardize rather than to say that it is a purely subjective guidance. There must be a difference in the guiding effect between the two. This requires us to design a simple and standardized flow chart to guide the bystander to implement CPR*.*”(P9)*

### 3.3 Theme three:Legal Factors

Relevant managers said that at present, there are no relevant laws in China to protect disputes encountered by dispatchers in telephone guidance.Dispatchers will be worried about telephone guidance, and their rights and interests will not be guaranteed fundamentally, so people are reluctant to implement T-CPR:

> *“I think the law is very necessary. “If there is no legal protection, the dispatchers are certainly worried about to implement T-CPR*.*Is it necessary for dispatcher to assume responsibility if he or she instructs bystanders to do CPR in case of problems*.*”(P4)*

### 3.4 Theme four:Guiding Factors

Different guiding methods, whether the guiding software is useful or easy to use will significantly affect the implementation of T-CPR. It is crucial for bystanders to implement T-CPR in the first time. In fact,the coexistence of various guiding methods can promote the implementation of T-CPR:

> *“I think if our country want to do this work, it should coexist in many ways which will be suitable for different age groups and be accepted by different age groups. Because most of the elderly do not have smart phones, they are easy to accept the dispatcher’s telephone guidance. Young and middle-aged people are more likely to use mobile APP or visual telephone. Mobile APP method can give someone the chance to learn CPR in their spare time. A variety of guidance methods can complement each other”. (P8)*

### 3.5 Theme five:Financial Factors

The amount of capital investment obviously affects the implementation of T-CPR. The software development of T-CPR, the allocation of instructors, the improvement of dispatching guidance system and the training of bystanders’ CPR skills all need sufficient financial support.

> *“T-CPR can not be carried out without the financial support of the government. The increase of personnel and the improvement of informationization need financial support. In fact, the price of developing such a system is relatively high*.*”(P1)*

## 4 DISCUSSION

Our study reveals that different cultural levels of bystanders will affect the effect of T-CPR. If the bystander does not understand the guidance, it is obviously impossible to implement T-CPR. In fact, it has been 33 years since China implemented free nine-year compulsory education in six-year primary schools and three-year junior middle schools. The Chinese people, especially the adults under the age of 60, already have the basic knowledge for implementing T-CPR. But in the telephone, especially in the traditional audio phone, it is difficult to understand how to do when the OHCA occurs. If we strengthen the training and propaganda of relevant knowledge, so that the bystanders have a certain basis, it is obviously easy to understand and accept and begin to implement T-CPR.For the training of CPR knowledge and skills, China has attached great importance under the leadership of the Red Cross Society of China after the May 12 Wenchuan earthquake in Sichuan province in 2008. However, compared with the huge population of nearly 1.4 billion in China, the proportion of training is still very low and uneven, ranging from 1.0% to 11.8%. There is a huge gap between China and the United States and other countries (Huijun et al.,2019). Therefore, on the one hand, we still need to continue to increase the training intensity and breadth of CPR, on the other hand, it is urgent to carry out T-CPR. Because T-CPR, after all, is guided by professionals and its role is obvious.Therefore, this study is proposed that strengthen the training of bystanders in cardiopulmonary resuscitation knowledge and skills, enhance their awareness of first aid, which will make the public accept T-CPR easily.

This study finds that Chinese people’s awareness of first aid is weak. People always feel that dangerous are far away from them. But once it happens, they often tend to be emotionally volatile. They dare not do CPR or don’t do CPR. They just ask the emergency center to send an ambulance as soon as possible. They ask the dispatcher not to say anything else. This is obviously related to the weak first aid knowledge of the public in China. Non-professionals have not yet realized the importance of “time is life” for SCA patients. These require our society give more propaganda. We can popularize the basic knowledge and skills of CPR through various ways, arouse the public’s awareness of first aid, create an atmosphere of first aid in the whole society, and form a good situation for ordinary people to sign up for CPR training voluntarily. Institutional organizations and enterprises require their employees to be trained initiatively. T-CPR should be introduced in CPR training. All of these will greatly facilitate the implementation of TCPR.Develop appropriate T-CPR guidance process, standardize the guidance procedure, work in a multi-pronged manner and coexist with various guidance methods, which is an effective way to promote the implementation of T-CPR in China.

The traditional T-CPR guidance is mainly dispatcher guidance. However, this study finds that in China, dispatchers are lack of professional skills and dispatchers in some first-aid centers have been overloaded with dispatching work. So it is difficult to carry out T-CPR. Some dispatchers of first-aid centers do not have medicalbackground and cannot perform T-CPR. In cities where T-CPR has been carried out, this study finds that their systems are imported from the United States. But Chinese national conditions are different from those of the United States, so it needs to be localized. This suggests that if T-CPR is to be carried out in China at this stage, for some first-aid centers in good conditions we can import advanced dispatching guidance system from abroad and then localize them, improve them to accord with Chinese national conditions, unify the training of first aid dispatchers, standardize the guiding procedures, and then carry out T-CPR guided by dispatchers. But in some first-aid centers where are lack of dispatchers or lack of professional skills of dispatchers, doctors or nurses should be involved in pre-hospital first aid and be trained for carrying out T-CPR (Seyed Bagheri et al.,2019).In addition, TCPR guidance software based on mobile phones can be developed. Studies have shown that mobile phone application software guidance is better than telephone guidance in identifying the first compression time and some parts of compression action after SCA (Dong et al.,2020;Plata et al.,2019).By installing guidance software in smartphones, it can provide guidance for the witnesses who are willing to perform chest compression. It also can avoid the embarrassment that the publics are willing to implement CPR but not only lack official guidance but also lack legal support. Finally, the implementation rate of pre-hospital CPR for patients with cardiac arrest will be improved. This study also suggests that the coexistence of various guidance methods can promote the implementation of T-CPR. But no matter which guidance method, unified, simple, standardized flow chart is needed. It is very important to make the bystanders understand it and implement T-CPR in the first time.Using internet technologies to build smart pre-hospital emergency system may promote the implementation of T-CPR.

At present, the world is in the information era of vigorous development of big data, cloud computing, artificial intelligence and other frontier technologies. Internet technology is being deeply integrated with all walks of life. The internet information has also become an important measure to raise the level of pre-hospital rescue. The early diagnosis of SCA and early positioning of SCA patients may be achieved by building a smart pre-hospital emergency system. This study offers some new insights on it. For example, if the wechat, which is widely used in China, can send the specific positioning of the patients to the dispatching center at the first time and provide the bystanders guidance of audio, video and text of CPR, so that more SCA patients will be treated. Smart pre-hospital emergency system is also possible to achieve intelligent calculation of the best route, so as to minimize emergency response time, and gradually achieve ‘accurate dispatch’ and ‘precise treatment’.

As this study shows that China has not yet enacted relevant laws at the national level to protect dispatchers from disputes encountered in telephone guidance. The dispatchers will have some misgivings because their rights and interests are fundamentally not guaranteed when they conduct telephone guidance. Previous studies by the research team also showed that a major concern of bystanders in implementing CPR is the lack of legal protection, especially when CPR is needed by the ‘strangers’ and ‘acquaintances’ (Xuehua et al.,2013). Therefore, it is urgent for the relevant departments to promote the legislation of first aid exemption as soon as possible, to clarify the scope and content of first aid exemption, so as to encourage more people to help patients with cardiac arrest and avoid tragedy. In recent years, Hangzhou, Zhejiang Province, China, began to implement the Regulations on Pre-hospital Emergency Management in Hangzhou, which put forward the public ‘first aid exemption’ and commendation regulations.However, it is necessary to further promote all cities and regions in China.Promoting the legislation of first aid exemption as soon as possible may promote the development of T-CPR.

Financial support from various channels, such as the state, local government or enterprises may promote the development of T-CPR.This study shows that the amount of investment has a significant effect on the implementation of T-CPR. The software development of T-CPR, the allocation of instructors, the improvement of dispatching guidance system and the training of bystanders’ CPR knowledge and skills all need sufficient financial support. Some economically developed cities, such as Beijing and Shanghai, can rely mainly on local financial input; some economically underdeveloped areas can be realized through state financial allocation; and other successful enterprises can give back social funds as a useful supplement. In short, life is more important than everything, and the value of life is paramount. We need to do more in first aid, which is also the most meaningful.

### 4.1 Limitations

Although this study adequately describes the experiences and opinions of some experts in the field of first aid in China, it is not without limitations.As with all qualitative research, the aim was to generate hypotheses and therefore lacks generalizability to all experts’ experiences and views in all areas of emergency care in China.The researchers acknowledge that most of the selected research objects are from developed cities in China, and the sample size is small. Due to regional limitations, the sample representativeness will be affected, so it should be improved in future studies.The coding of this study was conducted by one researcher, which presents both benefits and potential bias. Future studies could employ multiple coders to analyze interview content.

### 4.2 Conclusion

In order to save SCA patients more effectively, urge bystanders to implement CPR as soon as possible and ensure the quality of CPR implementation, it is urgent to promote the implementation of T-CPR in China. Through strengthening the training of bystanders’ knowledge and skills of cardiopulmonary resuscitation, raising the awareness of first aid in the whole society may provide a solid foundation for the implementation of T-CPR. Through the development of T-CPR guidance process suited to Chinese national conditions, standardization of guidance procedures, multi-pronged approach and coexistence of various guidance methods may provide an effective way for the implementation of T-CPR. Using internet technologies to build smart pre-hospital emergency system may provide technical support for the implementation of T-CPR. It is necessary to promote the legislation of first aid exemption as soon as possible to clear up the concerns for the implementation of T-CPR. It needs financial support from various channels, such as the state, local government or enterprises, to provide adequate guarantee for the implementation of T-CPR.

## Acknowledgements

This research was supported by the Basic Public Welfare Research Program of Zhejiang Province under Grant No.LGF20G030006.The authors would like to express thanks to the above sponsoring organizations,and all staff members for taking part in this research and for their openness.

## Funding Source

This work was supported by the Basic Public Welfare Research Program of Zhejiang Province [grant number No.LGF20G030006].

## Authors’ contributions

Study Design: Li Gui, Xuehua Zhu,Yin Lin.

Data Collection:Xuehua Zhu, Ying Chen,Yin Lin.

Data Analysis:Xuehua Zhu, Ying Chen,Yin Lin.

Manuscript Writing: Xuehua Zhu,Li Gui,Ying Chen.

## Notes

### Competing Interest Statement

The authors have declared no competing interest.

### Summary of Updates

Section on discussion updated to clarify limitations

